# The paradigm shift: heartbeat initiation without “the pacemaker cell”

**DOI:** 10.1101/2022.11.04.515227

**Authors:** Victor A. Maltsev, Michael D. Stern

**Author notes:** **Corresponding author:** Michael D. Stern.

## Abstract

The current dogma about the heartbeat origin is based on “the pacemaker cell”, a specialized cell residing in the sinoatrial node (SAN) that exhibits spontaneous diastolic depolarization triggering rhythmic action potentials (APs). Recent high-resolution imaging, however, demonstrated that Ca signals and APs in the SAN are heterogeneous, with many cells generating APs of different rates and rhythms or even remaining non-firing (dormant cells), i.e. generating only subthreshold signals. Here we numerically tested a hypothesis that a community of dormant cells can generate normal automaticity, i.e. “the pacemaker cell” is not required to initiate rhythmic cardiac impulses. Our model includes (i) non-excitable cells generating oscillatory local Ca releases and (ii) an excitable cell lacking automaticity. While each cell in isolation was not “the pacemaker cell”, the cell system generated rhythmic APs: the subthreshold signals of non-excitable cells were transformed into respective membrane potential oscillations via electrogenic Na/Ca exchange and further transferred and integrated (computed) by the excitable cells to reach its AP threshold, generating rhythmic pacemaking. Conclusions: Cardiac impulse is an emergent property of the SAN cellular network and can be initiated by cells lacking intrinsic automaticity. Cell heterogeneity, weak coupling, subthreshold signals, and their summation are critical properties of the new pacemaker mechanism, i.e cardiac pacemaker can operate via a signaling process basically similar to that of “temporal summation” happening in a neuron with input from multiple presynaptic cells. The new mechanism, however, does not refute the classical pacemaker cell-based mechanism: both mechanisms can co-exist and interact within SAN tissue.

## INTRODUCTION

During a normal life span the human heart executes more than 2 billion rhythmic contraction cycles, pumping blood to all organs commensurate with body demand at any given condition. Our understanding of the heartbeat origin has been evolving with the advances in technology, experimental techniques, ideas, and computation power. A major milestone in 1907 was the discovery of the sinoatrial node (SAN), a small piece of specialized heart tissue that paces heart contractions (Keith and Flack, 1907). In 1937, a slow potential change (i.e. the diastolic depolarization in current terminology) preceding the discharge of cardiac impulses was discovered by Arvanitaki working on the snail heart (Arvanitaki et al., 1937). Based on Hodgkin-Huxley formalism, Noble (Noble, 1962) explained the origin of the diastolic depolarization of “the pacemaker cell” as an interplay of kinetics of membrane ion currents. Further studies proposed a critical contribution of intracellular Ca cycling in addition to the membrane ion channels, giving rise a modern theory of the coupled-clock system (Maltsev and Lakatta, 2009) or ignition process via positive feedback mechanisms among local Ca releases (LCRs), Na/Ca exchanger (NCX) and L-type Ca channels (particularly of Cav1.3 isoform) (Lyashkov et al., 2018;Maltsev et al., 2022b).

At the SAN tissue level, it was initially thought that the pacemaker cell or a leading pacemaker center dictates the excitation rate and rhythm of other SAN cells (Sano et al., 1978;Bleeker et al., 1980). Subsequent studies suggested, however, that individual SAN cells mutually entrain each other to fire action potentials (APs) with a common period but different phases (dubbed “democratic” process) resulting in “apparent” propagation (Jalife, 1984;Michaels et al., 1987). Electrical mapping and imaging of SAN tissue demonstrated concentric excitation propagation from a leading center to SAN periphery (Efimov et al., 2004;Lang et al., 2011;Li et al., 2017). Thus, regardless of the propagation is real or apparent, the present dogma is that the pacemaker cell (possessing intrinsic automaticity) or a group of such cells with the fastest spontaneous diastolic depolarization rate within the SAN tissue paces or entrain other SAN cells and the entire heart.

While the spontaneously beating pacemaker cell(s) is thought to initiate cardiac impulse, only 10-30% of isolated cells actually contract spontaneously (Nakayama et al., 1984). The remaining nonfiring cells could not be ascribed as the true pacemaker cells and were not studied. Also, there has been always an uncertainty whether these cells were damaged by the enzymatic isolation procedure. Recent studies, however, demonstrated that this major population of non-firing cells (dubbed dormant cells) is not “dead”, but generates subthreshold oscillatory signals and many dormant cells can be reversibly “awakened” to generate normal automaticity in the presence of β-adrenergic receptor stimulation (Kim et al., 2018;Tsutsui et al., 2018;Tsutsui et al., 2021;Louradour et al., 2022). Dormant cells isolated from superior or inferior regions of SAN demonstrated different properties of noise of Ca signals and membrane potential (V_m_) subthreshold oscillations (Grainger et al., 2021), indicating that the role of these cells differs in different SAN locations.

While earlier low-resolution imaging of SAN tissue showed concentric continuous spread of excitation propagation, recent studies performed at a higher (individual cell) resolution level discovered a major fraction of non-firing cells (Bychkov et al., 2020;Fenske et al., 2020), and the tissue excitation within the SAN center appeared to be discontinuous, consisting of functional cell communities (presumably functional modules) with different firing patterns (Bychkov et al., 2020). Ca signals and APs in the communities are highly heterogeneous, with many cells generating APs of different rates and rhythms or remaining non-firing, generating subthreshold signals (Video 1). Thus, a possible functional role of non-firing cells and their subthreshold signals represents an open problem in cardiac biology.

One idea is that synchronized APs at the SAN exits emerge from heterogeneous subcellular subthreshold Ca signals, similar to multiscale complex processes of impulse generation within clusters of neurons in neuronal networks (Bychkov et al., 2020;Bychkov et al., 2022). Possible specific mechanisms of signal processing within the network also include stochastic resonance (Clancy and Santana, 2020;Grainger et al., 2021), a phase transition (Weiss and Qu, 2020;Maltsev et al., 2022a), and electrotonic interactions of firing cells with non-firing cells (Fenske et al., 2020).

Here we tested numerically a hypothesis that a fully functional pacemaker module generating rhythmic APs can be constructed exclusively from loosely connected dormant non-firing cells. Specifically, such module can be constructed with non-excitable cells (Ca oscillators) generating synchronized oscillatory local Ca releases driving rhythmic AP firing of an excitable dormant cell lacking intrinsic automaticity. Thus, cardiac impulse can be seen as an emergent property of the SAN cellular network/meshwork and it can be initiated by a community of dormant cells, i.e. without “the pacemaker cell” in its classical definition.

## METHODS

We used our previously published three-dimensional SAN cell model operating normally as a coupled clock system (Stern et al., 2014): (i) a Ca clock featuring individual stochastically operating ryanodine receptors (RyRs) embedded in the cell-distributed Ca store (sarcoplasmic reticulum) equipped with Ca pump and (ii) a membrane clock featuring the full set of electrophysiology equations describing V_m_ and 10 different ion currents (Maltsev et al., 2022b). The clocks are coupled via electrogenic Na/Ca exchanger and also via individual L-type Ca channels of two types, Cav1.2 and Cav1.3 differing in their activation voltages by 16 mV, stochastically operating and interacting with RyRs in couplons. We modelled a fully functional pacemaker module as a system of five loosely coupled dormant cells (Fig. 1A). Each dormant cell model was a modification of our original model to exclude spontaneous AP firing. Four cells in the pacemaker module were modelled as pure Ca oscillators (CaOsc cells), having relatively high sarcoplasmic reticulum Ca pumping rate P_up_ and Na/Ca exchange rate, but lacking I_CaL_ to prevent AP generation rendering cells non-excitable and non-firing, but generating subthreshold signals. These four CaOsc cells were connected to one cell that was modelled as a highly excitable dormant cell (AP cell) with large density of Cav1.2 but lacking automaticity in the absence of the low voltage activated Cav1.3. The idea for such functional module was inspired by our recent experimental observation in intact SAN tissue (Bychkov et al., 2022) that some SAN cells generate only oscillatory LCRs that precede AP-induced Ca transients in adjacent cells (Video 2). Thus, in our model the oscillatory subthreshold signals from the four CaOsc cells would be transferred via a low conductance connection to the membrane of the AP cell, with the AP cell membrane acting as an integrator (a computer) of the oscillatory signals that depolarize the cell membrane towards the activation I_CaL_ threshold, triggering an AP.

**Figure 1.**
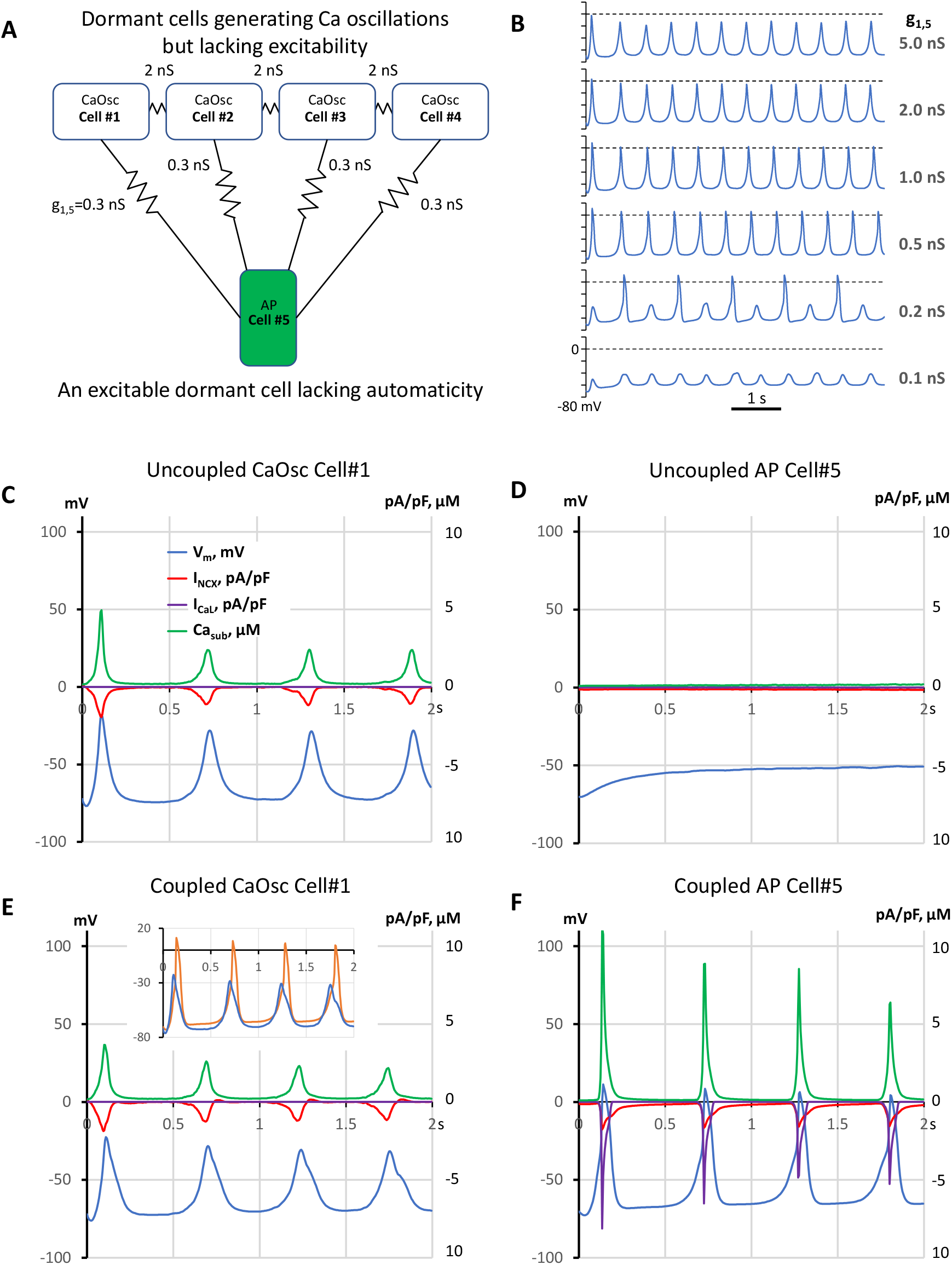
A model of emergent pacemaking in a network of 5 dormant cells (lacking intrinsic automaticity): A: A diagram of the network: cells #1-4 operating as Ca oscillators (CaOsc) and an excitable Cell #5 (i.e. AP capable cell, labelled as APC) are loosely connected (zigzag lines with conductance values). B: V_m_ time course of Cell #5 at various coupling with CaOsc cells (g_1,5_). C-F: Time series of I_NCX_, I_CaL_, V_m_, and mean submembrane Ca (Ca_sub_) simulated for CaOsc cell #1 and APC cell #5 in separation (C and D) and when cells were coupled (E and F) as shown in panel A. Inset in E shows overlapped V_m_ in coupled cells #1 (blue) and #5 (orange). V_m_ rise in cell #1 always preceded each AP upstroke in cell #5. See also related Video 3.

## RESULTS

First, we simulated each cell model behavior in separation (uncoupled cells). CaOsc Cell#1 generated oscillations of submembrane [Ca], NCX current (I_NCX_), and V_m_ (Fig. 1C,D). Importantly, the V_m_ oscillations were driven by respective I_NCX_ oscillations and had a relatively small amplitude (vs. an AP). At the same time, AP cell #5 alone failed to generate APs, with V_m_ fluctuating at a −50 mV level. Then we performed simulation for the coupled-cell system (Fig. 1, E,F). While the CaOsc cells #1-4 continued to generate subthreshold V_m_ oscillations, the AP cell #5 generated rhythmic APs and AP-induced Ca transients paced by cells #1-4. Importantly, the pacing signals always preceded each AP generation (inset in Fig.1E and Video 3). Next, we performed a parameter sensitivity analysis varying the coupling conductance between CaOsc cells and excitable AP cell (Fig. 1B). The model generated rhythmic APs in a wide range of coupling conductance from 0.3 nS to 1 nS. Finally, we performed simulations for a more complex network configuration with more cells (Fig. 2A) which demonstrated that CaOsc cells can drive more than just one AP cell (Cells #5 and #6) and the emergent automaticity can spread within further layers of dormant excitable cells (cells #7-9) (Fig. 2B).

**Figure 2.**
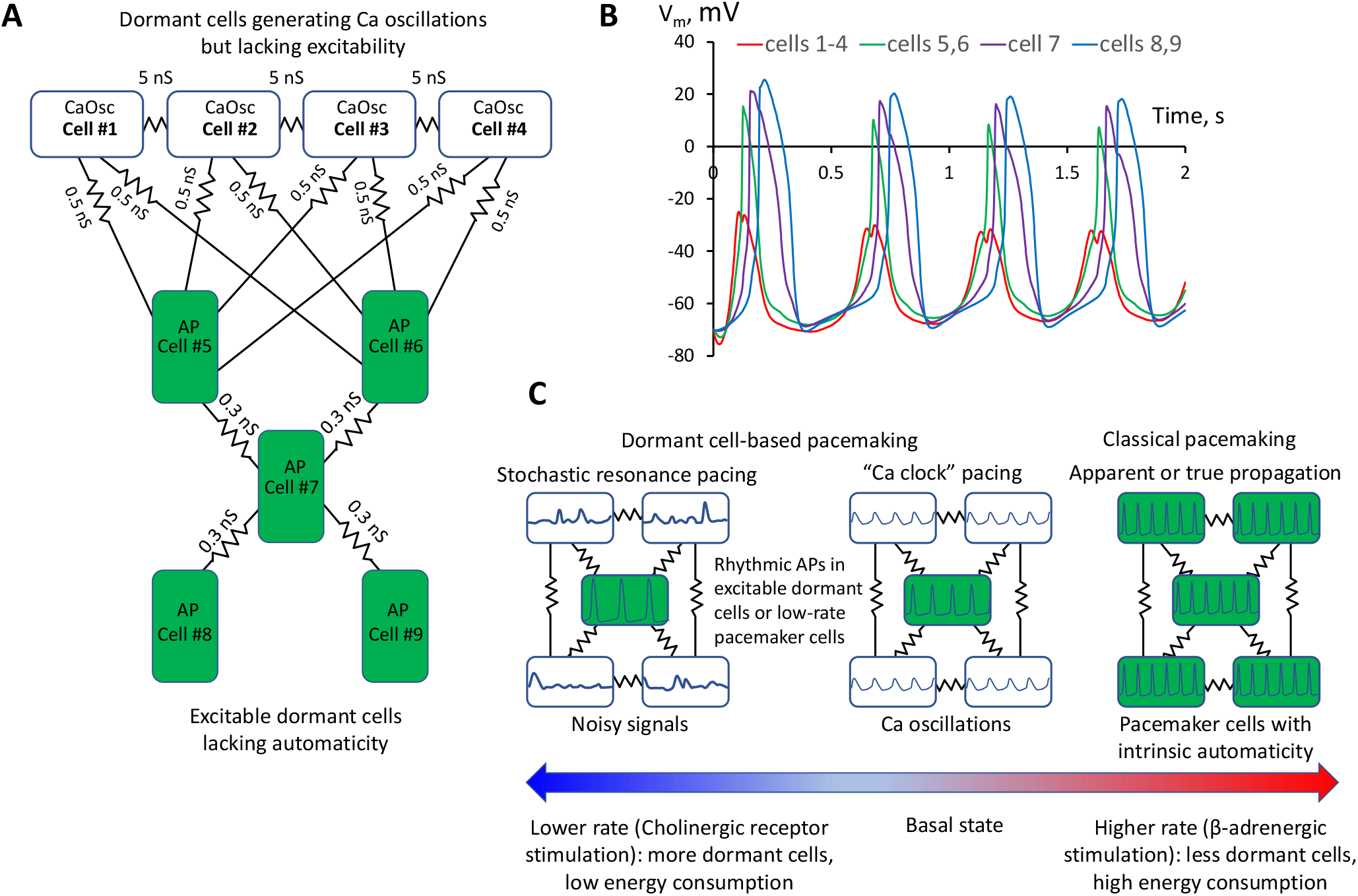
A more complex model of pacemaker network of 9 dormant cells demonstrating that Ca oscillator cells can drive more than one excitable cell and the emergent automaticity can spread within further layers of dormant excitable cells. A: A diagram of the network: cells #1-4 operating as Ca oscillators (CaOsc) and excitable Cells #5-9 (i.e. AP capable cells, labelled as APC) are loosely connected (zigzag lines with conductance values). B: V_m_ time course in the cells of the network. V_m_ rise in cell #1 always preceded AP upstrokes in excitable cells. APs in the 3 layers of excitable cells appear with respective delays reflecting interactions withing excitable cell layers. C: Prevalence of proposed pacemaking modes with respect to AP firing rates and autonomic modulation state: stochastic resonance pacing prevails at low rates and cholinergic receptor stimulation, but Ca clock pacing and pacemaker cell pacing prevail in basal state and during β-adrenergic stimulation.

## DISCUSSION

Here we demonstrate for the first time that rhythmic cardiac pacemaker impulses can be generated by an ensemble of heterogeneous (specialized) dormant SAN cells, i.e. cells lacking intrinsic automaticity. Such dormant cells have been previously identified in isolation (Kim et al., 2018;Tsutsui et al., 2018;Grainger et al., 2021;Tsutsui et al., 2021;Louradour et al., 2022) and in intact SAN tissue (Bychkov et al., 2020;Fenske et al., 2020). While the function of dormant cells remains unknown, it has been speculated that: 1) dormant cells stay “in reserve” and are “awaken” to function by β-adrenergic stimulation in the fight-or-flight response (Kim et al., 2018;Louradour et al., 2022); 2) non-firing cells can influence the frequency of AP firing cells via electrotonic interactions (Fenske et al., 2020). Experimental studies, however, have shown that dormant cells are not static or neutral: they do generate subthreshold signals (Kim et al., 2018;Bychkov et al., 2020;Grainger et al., 2021). One specific pattern of Ca signaling observed in experimental recordings in intact SAN tissue was that “some SAN cells generate only LCRs that precede AP-induced Ca transients in adjacent cells” (see an example in Video 2). It was further speculated that such subthreshold signals can somehow spatiotemporally summate and prompt initiation of full-scale signals (i.e. APs) within SAN tissue, e.g. via a stochastic resonance (Clancy and Santana, 2020;Grainger et al., 2021;Guarina et al., 2022), or a phase transition-like process each cycle (Weiss and Qu, 2020;Maltsev et al., 2022a). Thus, following this logic, in addition to the tonic influence or simple “wake-up” function under stress, dormant cells can be in fact functionally active generating important subthreshold signals. Here we suggest a simple model of multi-cellular pacemaker that can indeed operate via synchronization and integration of the subthreshold signals in a community of heterogeneous dormant cells that can reach a threshold each cycle to generate rhythmic APs and the excitation can spread in subsequent layers of dormant cells (Figs 1,2).

The requirement of heterogeneity for such synergistic pacemaker system in our model is in line with that SAN tissue indeed represents an extremely heterogenous structure (Boyett et al., 2000;Lyashkov et al., 2007). For example, major membrane cell conductances such as of I_CaL_ and I_f_ vary by an order of magnitude (Honjo et al., 1996;Monfredi et al., 2018) (Fig. S1). Dormant cells represent a major cell population (>50%) isolated from SAN (Nakayama et al., 1984;Kim et al., 2018;Grainger et al., 2021;Tsutsui et al., 2021;Louradour et al., 2022). Thus, the SAN tissue seems to be perfectly poised to such operation in the basal state and especially under parasympathetic stimulation, i.e. when the dormant cells become even more abundant in the SAN (Fenske et al., 2020).

Our model is robust: it generates rhythmic APs in a wide range of CaOsc-AP cell coupling conductance from 0.3 nS to 1 nS (Fig.1B,F). Weaker coupling generates alternations or subthreshold oscillations because the integrated impact of CaOsc cells is simply insufficient to bring V_m_ of AP cell to its AP threshold. Stronger coupling does bring V_m_ to the AP threshold, but AP cell cannot generate a full-scale AP because its electrotonic load of neighboring cells increases (Fig. 1B). Thus, another requirement of our model is weak cell coupling that allows cells to function independently, i.e. specialize as Ca clock or AP generator. This requirement is met in the SAN center which lacks high conductance CX43 gap junctions (Boyett et al., 2006;Bychkov et al., 2020).

How would the new pacemaker mechanism work in real SAN tissue? The real SAN tissue exhibits complex network configurations with substantial heterogeneity of the cell coupling strength and parameters of the interacting cells which, in turn, are controlled by autonomic modulation. Based on our results, in addition to pure dormant-cell based pacemaking, we can envision the existence of a transitional (mixed) type of network peacemaking, in which high-frequency Ca oscillators of dormant cells would pace cells featuring low-rate intrinsic automaticity. On the other hand, in some dormant cells Ca releases can be noisy and dysrhythmic (Kim et al., 2018;Grainger et al., 2021), i.e. not that large and rhythmic as in the model presented here. However, the concept of dormant cell-driving pacemaking proposed here may also work for such cells. An interesting possibility (to be tested in the future) is that the overlapping V_m_ fluctuations of these “noisy” cells can prompt AP generation in the neighboring cells (dormant cells or low-rate pacemaker cells) when they are ready to fire after a refractory period, or, in other terms, via stochastic resonance (Clancy and Santana, 2020;Grainger et al., 2021), when the noise components resonate with intrinsic frequency of AP generators. The 4-17 mV amplitude of noise reported in dormant SAN cells isolated form mouse (Grainger et al., 2021) was smaller than ~40 mV in our rabbit CaOsc cell model (Fig.1). V_m_ oscillation amplitudes can differ in dormant cells of different species and under different physiological and pathological conditions. For example, a wide range of V_m_ oscillation amplitudes (including those close to 40 mV in our model) was reported in isolated human and rabbit dormant SAN cells in their transition to AP firing state in response to β-adrenergic stimulation (Tsutsui et al., 2018;Tsutsui et al., 2021).

Hypothetical contribution of different pacing modes with respect to heart rate and its autonomic control is shown in Fig. 2C. Previous studies showed that various degrees of LCR synchronization are achieved (and controlled) by respective various degrees of CaMKII and PKA-dependent phosphorylation of Ca cycling proteins (Sirenko et al., 2013). At the basal state and especially during β-adrenergic response, higher levels of cAMP and PKA-dependent phosphorylation increase LCR synchronization (Vinogradova et al., 2006) that would favor Ca clock- and classical pacemaker cell-based pacemaking among cells in local neighborhoods in SAN tissue. The role of dormant cell-based pacemaking (and especially its stochastic resonance subtype) is expected to increase at low rates as cholinergic receptor stimulation increases the number of dormant cells (Fenske et al., 2020) and decreases LCR synchronization (Ca clock disintegrates, LCRs become noisy and smaller) due to lower cAMP production and attendant decrease in PKA-dependent phosphorylation of Ca cycling proteins (Lyashkov et al., 2009).

In addition to Ca clock (i.e. LCR synchronization), we also want to emphasize the roles of NCX and I_CaL_ that appear to be important not only for intrinsic automaticity of individual SAN cells (Maltsev and Lakatta, 2009;Lyashkov et al., 2018;Yue et al., 2020), but also in SAN tissue pacemaking: subthreshold Ca signals of synchronized LCRs are translated to respective V_m_ oscillations via NCX and further summated (i.e. computed) by the membrane of the AP generating cells in the neighborhood to bring their V_m_ to the threshold of I_CaL_ activation and generate AP upstroke. Therefore, the AP generating cell lacking automaticity in our model must have a substantial contribution of high-threshold activation Cav1.2 in order to be highly excitable but, at the same time, remain dormant. Increasing contribution of low-threshold activation isoform Cav1.3 initiates ignition process that drives automaticity (Torrente et al., 2016;Lyashkov et al., 2018;Maltsev et al., 2022b). Thus, the amplitude of I_CaL_ and its fraction of Cav1.2/Cav1.3 components may also regulate the occurrence and contribution of different modes of pacemaking in cell communities.

What are potential roles and benefits of the new pacemaker mechanism? It can keep the system alive at the edge of criticality in heterogenous SAN tissue (Campana et al., 2022;Maltsev et al., 2022a), e.g. in aging and disease. Indeed, major pacemaker components (I_CaL_, I_f_, and LCRs) of intrinsic automaticity deteriorate with aging (Jones et al., 2007;Larson et al., 2013;Liu et al., 2014). Thus, in the absence of classical pacemaker cells, the dormant cell-based pacemaker mechanism might serve as the last resort keeping the heart ticking. Further, it can save energy. Indeed, dormant cells, generating subthreshold signals, do not generate high energy-demanding APs. During fight-or-flight response (critical for survival) the dormant cells “awake” (Kim et al., 2018;Tsutsui et al., 2018;Tsutsui et al., 2021;Louradour et al., 2022) and system operation can shift the gear from its energy-saving mode to classical paradigm of pacemaking driven by classical pacemaker cells at higher rates (Fig. 2C).

In more general context, the shift of paradigm from pacemaker-cell based to network-based type of pacing is not unique: a similar paradigm shift has happened recently in studies of respiratory rhythm: the respiratory oscillations arise from complex and dynamic molecular, synaptic and neuronal interactions within a diverse neural microcircuit, and “pacemaker properties are not necessary for respiratory rhythmogenesis” (Feldman and Kam, 2015). However, it is important to emphasize that the new dormant cell-based pacemaker mechanism does not refute or substitute the classical pacemaker cell-based mechanism: both mechanisms can co-exist and synergistically interact in the form of different functional modules within the central SAN area (Bychkov et al., 2020;Norris and Maltsev, 2022) with different prevalence in different conditions (Fig. 2C). Rather, our results challenge the exclusive role of the fully automatic pacemaker cells in heart pacemaker function, because a fully functional pacemaker can emerge from subthreshold signal integration (computation) within a dormant cell community.

Elucidation of the role of the new mechanism will need more experimental and theoretical studies. It remains to be determined how different mechanisms of network automaticity would shift and operate in the presence of various densities of ion currents (I_f_, Ca_v1.2_ and Ca_v1.3_, I_CaT_, I_Kr_, I_NCX_, I_KACh_, etc.) and Ca clock activities (SR Ca pump and RyRs) in models with larger cell numbers featuring more complex (realistic) cell network structure, coupling strength, and autonomic modulation state, i.e. various levels of cAMP and CaMKII- and PKA-dependent phosphorylation of both Ca- and membrane-clock proteins.

The numerical prototype of dormant cell-based pacemaker presented here is based on our previous single cell model (Stern et al., 2014) that simulates transitions of each RyR and L-type Ca channel molecule and diffusion among numerous voxels in the entire network SR and cytosol (including natural buffers) in 3 dimensions. While this specific cell model generates detailed local Ca dynamics, including oscillatory synchronized LCRs critical for the emergent pacemaking (Video 3), it has a high computational demand. Future studies of SAN pacemaker mechanisms (including dormant-cell based pacemaking) in larger tissue models will likely use simpler cell models in the spirit of multi-scale modeling approach (Qu et al., 2011), such as common pool models featuring coupled-clock function (Campana et al., 2022;Maltsev et al., 2022a) or CRU-based models, generating LCRs (Maltsev et al., 2022b).

## FUNDING

This research was supported by the Intramural Research Program of the National Institutes of Health, National Institute on Aging

## ACKNOWLEDGEMENTS

This study utilized the high-performance computational capabilities of the Biowulf Linux cluster at the National Institutes of Health, Bethesda, MD. (https://hpc.nih.gov/)

## CONFLICT OF INTEREST

The authors declare that the research was conducted in the absence of any commercial or financial relationships that could be construed as a potential conflict of interest.

## SUPPLEMENTARY MATERIAL

It includes 3 videos and one supplementary figure (Fig. S1)

### Video legends

**Video 1**

An example of heterogeneous Ca signaling in central mouse SAN that includes rhythmic generation of AP-induced Ca transients in some cells and local subthreshold signals (LCRs) in other cells. Ca recording from (Bychkov et al., 2020).

**Video 2**

An example of a local Ca signaling module in central mouse SAN in which some SAN cells generate only LCRs that precede AP-induced Ca transients in adjacent cells. Ca recording from (Bychkov et al., 2020). For exact anatomical location within intact SAN and detailed analysis of this video, see Fig. 6 in (Bychkov et al., 2020).

**Video 3**

Simulation results of our new pacemaker cell-coupled system comprised of dormant cells (Fig. 1). Upper panel: Oscillatory synchronized LCRs under cell membrane in non-excitable Cell #1 (Ca oscillator). Lower panel: AP-induced Ca transients in excitable Cell #5, lacking intrinsic automaticity but paced by Ca oscillators in cells #1-5. Simulation duration is 2 s (see related time course of Vm, Fig. 2C,D). Note: Ca oscillations in cell #1 always precede AP-induced Ca transients (global Ca spike) in cell #5. The panels are the cell cylinder surfaces unwrapping to squares with distributions of RyR clusters under the cell membrane shown by cyan shades reflecting Ca loading of the junctional sarcoplasmic reticulum in which the clusters are embedded: 0 is pure black and >600 μM is pure cyan. Local Ca dynamics is shown by red shades: 0.1 μM is pure black (resting Ca level) and >5 μM is pure red. Simulations were performed using our modified model of SAN cell in 3 dimensions with individual RyRs and L-type Ca channels (Stern et al., 2014).

## Notes

### Competing Interest Statement

The authors have declared no competing interest.

### Summary of Updates

1. We performed sensitivity analysis for connectivity conductance (new Figure 1B) to illustrate importance of weak coupling 2. We report results with a more complex model with 4 layers of dormant cells (new Figure 2) to illustrate that the subpopulation of CaOsc cells can drive more than just one excitable cell and the emergent automaticity can spread within further layers of dormant excitable cells. 3. We added new supplemental figure S1 with our previously published patch clamp data demonstrating substantial cell-to-cell variability of ICaL and If. 4. We added a new diagram (Figure 2C) illustrating possible importance (prevalence) and shifts between different pacing modes in different conditions.

