## Supplementary material for "The paradigm shift: heartbeat initiation without “the pacemaker cell”": Fig. S1

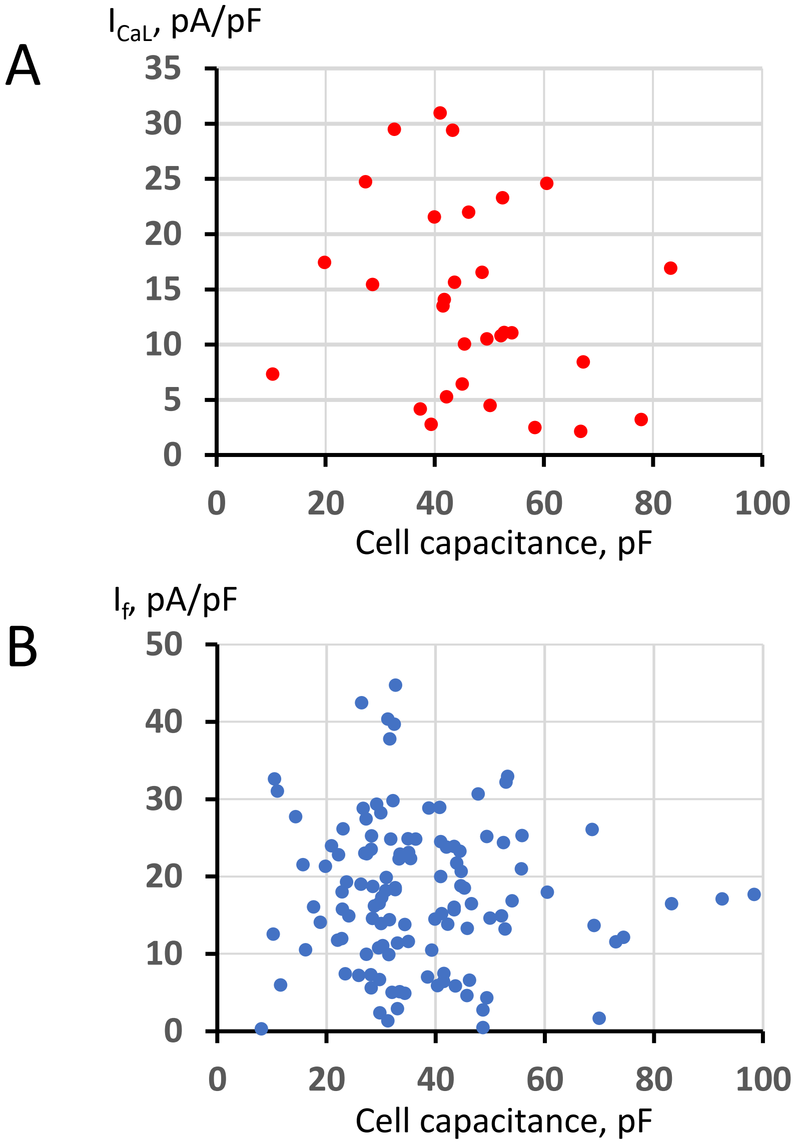


**Figure S1**. Substantial cell-to-cell variability of densities of L-type current ($I_{CaL}$, panel A) and funny current ($I_f$, panel B), measured by whole-cell patch clamp technique in isolated rabbit SA node cells of different membrane capacitance (i.e. different sizes). Modified from Monfredi, O., Tsutsui, K., Ziman, B., Stern, M.D., Lakatta, E.G., and Maltsev, V.A. (2018). Electrophysiological heterogeneity of pacemaker cells in the rabbit intercaval region, including the SA node: insights from recording multiple ion currents in each cell. Am J Physiol Heart Circ Physiol 314, H403-H414. doi: 10.1152/ajpheart.00253.2016.
